# Agricultural pressures impair trophic link between aquatic microorganisms and invertebrates

**DOI:** 10.1101/2021.03.18.435985

**Authors:** Rody Blom, S. Henrik Barmentlo, Maarten J.J. Schrama, Ellard R. Hunting

## Abstract

Decadal declines in aquatic ecosystem health prompted monitoring efforts and studies on effects of human practices on aquatic biodiversity, yet a consideration of ecological processes and trophic linkages is increasingly required to develop an in-depth understanding of aquatic food webs and its vulnerability to human activities. Here, we test in laboratory incubations using natural organic matter whether agricultural practices have an effect on two interacting ecological processes (i.e., decomposition and invertebrate growth) as the relevant temporal components of the trophic linkage between aquatic microbial communities and aquatic invertebrates. We further assess whether these altered trophic interactions are visible on ecologically relevant scales. We observed clear patterns in agricultural constraints on microbial decomposition, which coincided with reduced invertebrate growth and an unexpected increase in invertebrate consumption of organic matter. Similar differences in invertebrate length depending on land use were observed in our field survey, thereby providing important clues on the relevance and vulnerability of interdependent processes that can serve to improve future forays in monitoring ecosystem health.

## Introduction

Globally, anthropogenic activities pose significant threats to aquatic ecosystems (Strayer & Dudgeon, 2010; Vörösmarty et al., 2010; Strayer, 2010a). Resulting declines in ecosystem health have fueled monitoring activities, which are primarily aimed at quantifying biodiversity and ecosystem processes to allow for diagnosing perturbations and safeguarding the natural environment. A key ecosystem process proven vulnerable to such stressors is the degradation of dead organic matter (OM), such as decaying plant litter, that serves as food for aquatic microorganisms and invertebrates (Webster & Benfield, 1986; Wallace & Webster, 1996; Bundschuh & McKie, 2016). This process relies on a complex interplay between microorganisms and invertebrates where microbial conditioning of OM enhances nutritional value for invertebrates, while consumption of OM by invertebrates renders a diverse size range of dissolved and particulate OM that supports microorganism as well as filter-and deposit feeding invertebrates (Barlöcher, 1985; Graça et al., 1993; Graça, 2001; Wright & Covich 2005; Danger et al., 2012; Danger et al., 2012; Arce Funck et al., 2015; Vonk et al., 2016; Bundschuh & McKie, 2016). While most studies and monitoring efforts focus on diversity measures such as species abundances within a single trophic level, an explicit consideration of the complexity of interactions is now crucial to develop an in-depth understanding of aquatic food webs and its vulnerability to human activities (Hines et al., 2016; Schrama et al., 2017; Seibold et al., 2018; Bruder et al., 2019).

Agricultural practices are a primary source of pollution to aquatic sytems. This is due to the use of chemicals such as pesticides and fertilizer that run-off to adjacent aquatic systems where they can have direct- or indirect toxic effects in the water column or associate with OM accumulating in their sediments (e.g., Knezovich et al., 1987; Vijver et al., 2017). To date, studies aimed at assessing effects of agricultural practices mostly focussed on direct effects (e.g., toxicity) to aquatic invertebrates, while studies focused on indirect effects of chemicals on invertebrates relied predominantly on feeding preference experiments and singled-out chemicals that are often applied to the overlying water (e.g. Feckler et al., 2016; Vijver et al., 2017; Rossi et al., 2018; Barmentlo et al., 2018, 2019). This approach prevents the distinction between direct and indirect effects on microbially-mediated trophic linkages that are caused by chemical deposition and sorption of chemical to OM on temporal scales that are relevant to ecosystem processes.

Since various chemicals have been observed to affect microbial community composition and diversity (Gardeström et al., 2016; Tlili et al., 2016; Hunting et al., 2017a, Fasching et al., 2019), alterations in microbial diversity can be expected to affect the overall rate of microbial decomposition. This is relevant since the process of microbial decomposition, in contrast to microbial diversity, provides a more reliable reflection of OM-conditioning and available microbial biomass, and therewith the resources available for invertebrates, for the time period (seasons to years) that is required for OM to decompose and invertebrates to develop (e.g., Hines et al., 2016). However, as with invertebrates, the vast majority of studies on microbial communities rely on single time point characterization of microbial diversity or exposure to manipulated concentrations of model chemical compounds. It thus remains uncertain whether complex mixtures of organic particles and pollutants currently present in the environment translate to altered microbial processes, and whether changes in the temporal attributes of microbial processing of OM affect invertebrate performance.

This study tests whether the chemical complexity associated with agricultural practices affects microbial processing of OM and if this translates to aquatic invertebrates that rely on microbial biomass and microbial conditioning of OM. To this end, we assessed the effects of agricultural practices on 1) the contribution of microbial communities on decomposition of OM derived from an agricultural area, and 2) how changes in OM microbial decomposition affect the growth of the aquatic invertebrate *Asellus aquaticus* under controlled laboratory conditions. Finally, we surveyed *A. aquaticus*’ body length in relation to agricultural practices within the same agricultural area.

## Material and methods

### Experimental approach and research setting

We performed both a laboratory feeding experiment and a field survey to determine the effect of agricultural practices on microbial OM decomposition and invertebrate growth. In both components of the study, we exposed *Asellus aquaticus* individuals to OM-rich sediments originating from ditches adjacent to the three key types of land use present in an agricultural area, ‘Bollenstreek’, the Netherlands (Lat: 52°15’55.66”, Long: 4°28’27.94”), that have well-documented effects on microbial community structure and decomposition (Hunting et al., 2016; 2017a). Land uses selected were a dune nature reserve, agricultural sites growing flower bulb (hyacinths, lilies, tulips and daffodils) and permanent grassland. The agricultural area is intensively used to grow flower bulbs, which are treated with pesticides and nutrients between February to November. The ditches are adjacent to the flower bulb fields, which results in the transfer of pesticides and nutrients from the fields into the ditches. The grassland pastures are used for grazing by livestock. Natural sandy dunes can be found to the north and north-west of the area at approximately 200m from the ditches. The water bodies located in this area are hydrologically isolated from the agricultural area because of a natural elevation gradient (Ieromina et al., 2015). All water bodies are characterized by a wide variety of aquatic invertebrates (Ieromina et al., 2015; Hunting et al., 2016).

### Feeding experiment

To assess whether changes in microbial decomposition of OM affect the growth of aquatic invertebrates, we performed a laboratory feeding assay where we placed *Asellus aquaticus* (being a common detritivore in many freshwater habitats) on a diet of OM collected from ditches adjacent to either flower bulb fields, grasslands or pristine dune area. *A. aquaticus* was chosen as a model organism as it is a highly abundant detritivore in aquatic ditch systems. We prepared Decomposition and Consumption Tablets (DECOTABs) as standardized OM substrate as described by Kampfraath et al. (2012) and Van der Lee (2020). In brief, DECOTABs (ø17 mm) were prepared from naturally-derived OM embedded in an agar matrix, containing OM derived from sediments adjacent to flower bulb fields, grasslands or pristine dune area. DECOTABs were prepared as explained in detail in Hunting et al., 2016. Individuals of *A. aquaticus* used in the experiment were reared in the laboratory. Prior to the experiment, an egg-carrying female was isolated and placed in Elendt M4 medium (OECD, 2004) on a small layer (~1 cm) of fine quartz sand. In total, 36 neonates of *A. aquaticus* were collected directly after hatching. As such, all collected neonates were from the same spawn which limits variation in body length between individuals at the start of the experiment. Subsequently, neonates were placed in 50 mL glass bottles individually, which were filled with 35 mL Elendt M4 medium. One DECOTAB was added to each bottle. In total, 36 bottles containing one *A. aquaticus* neonate and one DECOTAB each (n = 12 per land use type) were used in the experiment. Additionally, per treatment, three control bottles containing one DECOTAB and without invertebrates were used to assess substrate dependent microbial decomposition rates. The experiment was conducted at a constant temperature of 18 °C and a light-dark regime of 16:8 hours for the duration of 42 days. Elendt M4 medium was replaced with freshly prepared medium every seven days. Physicochemical parameters (pH, dissolved oxygen, electrical conductivity, and temperature) were measured weekly using a HACH HQ40D electronic multi meter to ensure they were comparable between treatments throughout the experiment (data not shown). From t=14 onwards, *A. aquaticus* individuals were photographed every seven days using an eScope DP-M17 USB-microscope camera to monitor growth rates. Subsequently, *A. aquaticus* body length was quantified by measuring from cephalothorax to pleotelson from a dorsal perspective using ImageJ (v1.47). Potential effects of DECOTABs per land use, time (in days), and their respective interaction on *A. aquaticus*’ body length were analyzed using a linear mixed effect model in R (R Core Team, 2019), with the individual animals added as random variable (to account for the repeated measures design). To meet the assumption of normality of the model residuals and the random variable, body length data was log10 transformed. Furthermore, analysis of mortality data was performed by means of a logistic regression. After termination of the experiment (42 days after inoculation), remaining DECOTAB material was collected, dried at 60 °C for 48 hours and weighed on a balance (BP210S, Sartorius AG). In addition, DECOTABs which were only exposed to substrate-dependent microbial decomposition were dried and weighed following the same protocol. In order to assess initial DECOTAB weight, ten unused DECOTABs per treatment were dried and weighed following the same. Thereafter, both microbial decomposition and isopod consumption of OM was analyzed using a one way-ANOVA with a Tukey HSD post-hoc test.

### Field survey

To determine the effect of agricultural practices on aquatic invertebrate growth in the field, we collected *A. aquaticus* individuals from the top sediment layer by grab sampling using a 500μm dipping net at the same sites where the organic material for the lab experiment was retrieved. All specimens were collected on the same day in June 2016. At each of the three sites corresponding with the three types of land use (bulb, grassland, and pristine dune area), roughly 20 *A. aquaticus* individuals were collected, with a total of 57 individuals. Sexes were determined by examining the morphological properties as described by Bertin & Cézilly (2003). Individual *A. aquaticus* collected from the field were photographed and measured as stated earlier. An analysis of the effect of land use adjacent to the habitat on the body length of field-collected *A. aquaticus* was conducted using a one-way ANOVA model with length of each specimen as a dependent variable and land use as a predictor. Because *A. aquaticus* sexes typically differ in body length, we carried out separate analyses for both sexes.

Agricultural practices in the study area have well documented effects on microbial decomposition and community composition (Hunting et al., 2016; 2017a). A full assessment of microbial parameters was therefore considered to be superfluous and beyond the scope of this study. To confirm that agricultural practices in the study area differentially affected microbial communities during this study period, we measured carbon utilization profiles of sediment bacteria as a proxy of microbial functional (metabolic) diversity using Ecoplates (Biolog). Ecoplates are comprised of ecologically relevant, structurally diverse compounds, yet do not include e.g., recalcitrant substrates nor specific substrates typical of the soils used in this study. It is therefore impossible to directly relate substrate utilization profiles to the actual functioning of the soil microbial communities. Nonetheless, the number of substrates used can serve as a proxy of the metabolic diversity of the microbial community (Garland, 1999). To this end, microbial communities were sampled from sediments adjacent to dunes, grasslands, and bulb fields (6 replicates per treatment). One mL of sediment was diluted 50 times with demineralized water and vortexed. Mineral substrate was allowed to settle and subsequently distributed over well plates (Biolog, Ecoplate) within 2 hours after sampling. Plates were incubated for 96 h at 18°C and absorbance was measured using a standard plate reader. Carbon utilization profiles were analyzed using principal component analysis and a Jaccard-based one-way analysis of similarity (ANOSIM).

## Results

After 42 days of incubation, *Asellus aquaticus* contributed significantly (ANOVA, F = 56.8, p < 0.001) more to DECOTAB mass loss, when compared to DECOTAB exposed solely to microbial biofilms (Fig 1A). A shift in relative contributions to OM degradation becomes evident when plotting the ratio of microbial decomposition and isopod-associated OM consumption (Fig 1B), illustrating a shift towards a higher contribution of microbial decomposition to OM degradation. Isopods contributed to 37.8% of biomass loss of dune-derived OM, whereas they contributed to 85.2% and 90.3% of biomass loss in bulb field- and grassland-derived OM, respectively. We observed major differences between the ratios of microbial decomposition and isopod-associated degradation of bulb field- and grassland derived OM on the one hand and dune-derived OM on the other hand.

**Figure 1.**
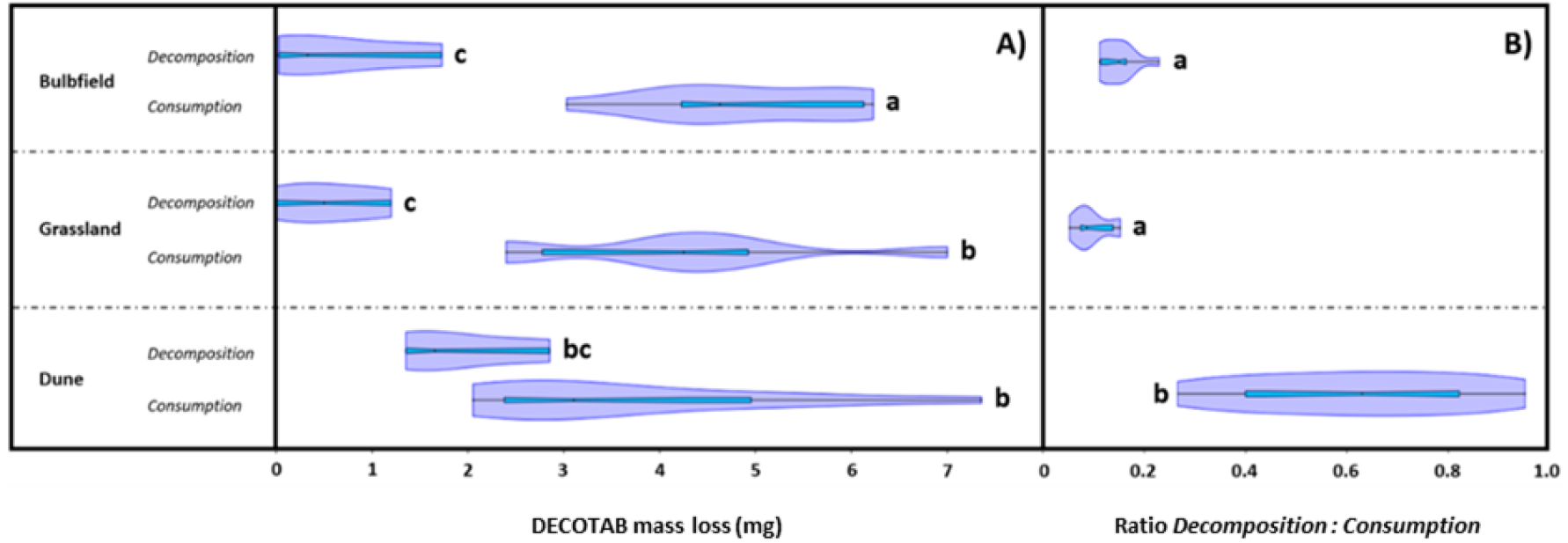
Organic matter mass loss after 42 days of microbial decomposition (± SE, n = 3 for dune, grass and bulb) for each land use type (ANOVA; significance level p = 0.05) and *Asellus aquaticus* consumption (A; ± SE, n = 12 for dune, n = 8 for grass, and n = 11 for bulb; only surviving individuals are shown) and ratio between microbial decomposition and isopod consumption of organic matter (B).

Over the course of 42 days, we observed that *A. aquaticus* feeding on OM derived from the dune area had significantly higher growth rates compared to *A. aquaticus* that fed on DECOTABs with grass- or bulb OM (TukeyHSD: p_dune-bulb_ < 0.001 and p_dune-grass_ < 0.001), whereas no difference in growth rates was observed between *A. aquaticus* that fed on bulb-and grass OM (TukeyHSD: p = 0.377; Fig. 2). In addition, *A. aquaticus* that fed on grassland-derived OM had lower survival (66.7%) at the end of the experiment compared to juveniles that fed on dune-(100%) and/or bulb (91.7%) derived OM (Chi-square, p = 0.03).

**Figure 2.**
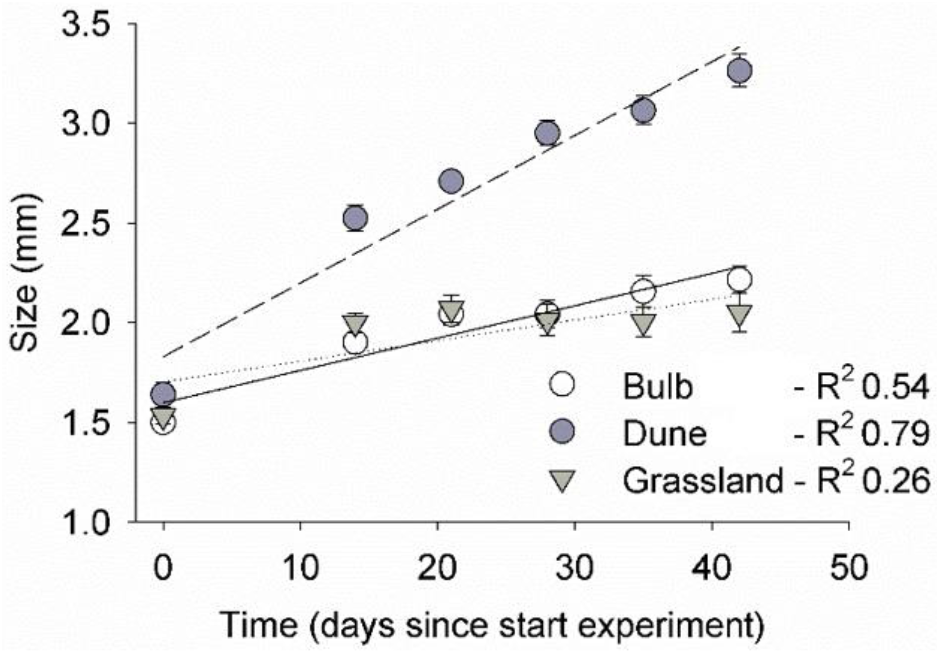
Body length (± SE, n = 12) of *Asellus aquaticus* during 42 days of incubation feeding on DECOTABs inoculated with organic matter from ditches adjacent to bulb fields (y = 1.69 + 0.011x), pristine dune area (y =1.82+0.037x) grassland (y = 1.59 + 0.0163x) (LME, significance level p = 0.05). On average, *A. aquaticus* which fed on organic matter collected from ditches adjacent to pristine dune areas displayed a significantly higher growth rate in comparison to those which fed on organic matter collected from ditches adjacent to bulb fields and grasslands.

In our field survey, we observed differences in microbial carbon utilization profiles depending on land use (Fig. 3A: ANOSIM; p < 0.05). This indicates differences in metabolic- and functional diversity of microbial communities between land use types. Analysis of body length of both sexes combined suggests an effect of land use, as individuals collected from the dune area were larger than those collected from the bulb- and grassland area (LME, p = 0.042). Males constituted the majority of the collected *A. aquaticus* individuals were (40 males vs 17 females). Male *A. aquaticu*s were larger than female individuals (figure 3BC), which is commonly observed for this species (Bertin & Cézilly, 2003). Body length of *A. aquaticus* collected from the dune area were largest, and those from the grassland area were the smallest; individuals from bulb fields were of intermediate length and did not appear to differ from individuals from other land uses. This pattern was only observed in male individuals (Fig. 3B: TukeyHSD: p_dune vs grass_ < 0.001, p_grass vs bulb_ = 0.03). Female individuals collected from grassland ditches were smaller compared to individuals collected from ditches in dune areas (Fig. 3C: TukeyHSD: p_dune-grass_ = 0.049).

**Figure 3.**
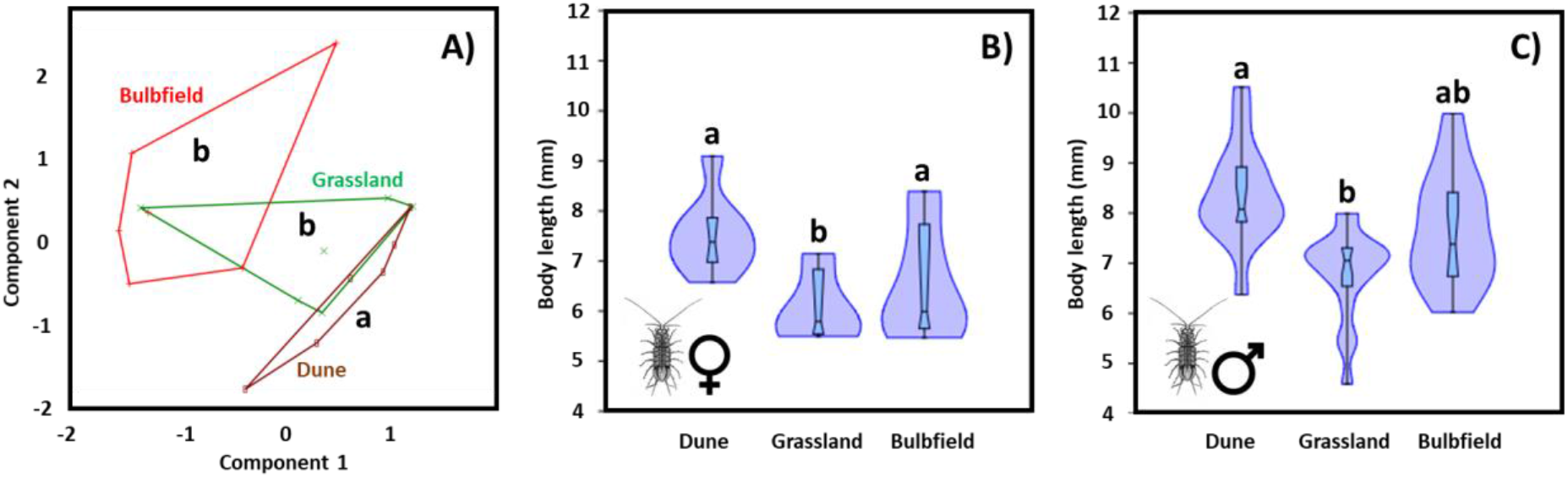
Microbial carbon utilization profiles of microbial communities extracted from sediments collected from ditches adjacent to dune-, bulb field- and grassland areas (A; n = 6; ANOSIM; p < 0.05) and average body length (± SE, B: n = 5-8; C: n = 14-18; ANOVA + Tukey post-hoc; significance level p = 0.05) of female (B) and male (C) *Asellus aquaticus* collected from ditches adjacent to a pristine dune area, grasslands and bulb fields. Corresponding letters indicate statistical similarity.

## Discussion

This study tested whether agricultural practices have an effect on two ecological processes (i.e., decomposition and invertebrate growth) that represent relevant temporal components of the trophic linkage between aquatic microbial communities and aquatic invertebrates, and assessed whether potential alterations are visible on ecologically relevant scales. We observed clear patterns in agricultural practices on microbial decomposition of OM collected in the field, which cascaded to effects on higher trophic levels by reduced invertebrate growth of both laboratory incubated and field-collected invertebrates. This carries implications for a range of ecological settings and current assessments of ecosystem health.

While microbial communities in natural systems are often considered resilient to natural disturbances and species loss due to a certain level of functional redundancy (e.g., Langenheder et al., 2005), a wide array of chemical pressures (e.g., agricultural chemicals and nanoparticles) have been shown to govern microbial community assembly and decomposition (Tlili et al., 2016; Hunting et al., 2016; 2017a; Zhai et al. 2018;). Here, we also observed microbial communities in the field to differ in their metabolic potential, and OM collected in the field to differ in microbial decomposition depending on the type of agricultural practices. In addition to direct toxic effects of chemicals, different organic practices result in differences in OM-subsidy (e.g., different crops) as well as sorption of agricultural chemicals to OM, affecting both quantity and quality of OM. This is relevant as microbial community assembly and processes are frequently observed to be driven by quality of the available OM resources (e.g., Myers et al., 2001; Docherty et al.,2006; Strickland et al., 2009; Hunting et al., 2013). The observed adverse effects of agricultural chemical on microbial decomposition of OM may be relevant to a wide array of chemicals present in agricultural catchments, provided hydrophobic fungicides can also inhibit fungal hyphomycete growth and microbial community composition (Zubrod et al., 2015; Konschak et al., 2019).

In contrast to measures of microbial diversity, decomposition rates provide a more reliable reflection of the available microbial biomass over extended periods of time. To assess whether differences in microbial decomposition translate to differential performances in a higher trophic level, growth rates of *Asellus aquaticus* were measured in relation to OM decomposition. We observed that *A. aquaticus* performed better when feeding on OM derived from dunes compared to bulb- and grassland-derived OM. This suggests invertebrate growth follows similar patterns as microbial decomposition depending on type of land use, i.e., higher growth rates with higher microbial decomposition rates. Dissimilarities in palatability and nutritional value between the different OM sources and their associated microbial communities can thus become visibly in invertebrate growth and size. While direct effects of chemicals and their metabolites present in the OM on invertebrate performance cannot be excluded (Zubrod et al., 2010), reduced growth rates have been causally linked to an absence of OM-associated microorganisms, emphasizing the importance of microorganisms in the diet of invertebrate shredders (Zhai et al. 2018). This corroborates the notion that colonization and conditioning of OM by microorganisms plays an important role in invertebrate nutrition and invertebrate-mediated consumption of OM (Graça et al., 1993; Graça, 2001; Wright & Covich 2005; Danger et al., 2012; Hines et al., 2016), and suggests that observed growth reductions in response to agricultural practices is due to a decrease in microbial biomass availability over the course of the experiment. This seems to have prompted *A. aquaticus* to feed directly on OM rather than microbial biomass, as we observed an unexpected increase in consumption of bulb- and grassland-derived OM that apparently contained fewer essential nutrients such as fatty acids and proteins to support growth. This points to a currently underappreciated compensatory feeding mechanism in invertebrates in response to microbial perturbations.

Conventional monitoring efforts aimed at assessing ecosystem health generally relies on single time point quantification of species diversity and abundance with a large emphasis on invertebrates (Hunting et al., 2017b). Often, stressor-induced effects are not directly reflected by species diversity and abundance alone, but also by characteristics related to species-specific life-history, such as body length, that capture the long-term result of exposure to stress (Culp et al., 2010; Hines et al., 2016). Here, agricultural practices were observed to have an effect on two ecological processes (i.e., decomposition and invertebrate growth) that capture the temporal complexity of the trophic linkage between aquatic microbial communities and aquatic invertebrates. The observed adverse effects of agricultural stressors on the performance of OM-associated microbial communities and invertebrates likely ripples through the food web as it forms a key source of fine particulate OM for filter- and deposit feeding invertebrates (Bundschuh & McKie, 2016). Impaired invertebrate growth rates may also lead to reduced biomass available for predators that rely on invertebrate shredders as food source, such as fish (Rask & Hiisivuori, 1985) and invertebrate predators (Herrmann, 1984; Krisp & Maier, 2005). In other words, current monitoring efforts that rely on single time point estimation of invertebrate abundances fail to capture the temporal nature and complexity of trophic interactions (e.g., knock-on effects on growth rate and other fitness variables) that ultimately govern the functioning and health of ecosystems. Thus, the likelihood that monitoring efforts can lead to misinterpretations of ecosystem health assessments calls for reconsidering our approaches in biodiversity assessments and environmental diagnostics.

## Conclusion

Using OM that encapsulates the naturally relevant complexity of agricultural chemicals entering adjacent water bodies, this study shows that agricultural practices can reduce the contribution of microbial communities to OM degradation, which can coincide with a reduction in growth rates of the aquatic invertebrate *A. aquaticus* as assessed under controlled laboratory conditions. Quantifying length of *A. aquaticus* in the field in ditches adjacent to different agricultural practices reveals that the patterns observed in our laboratory study reflect those occurring in natural systems. In a context of natural complexity and interdependence of ecological processes, these results suggest that agricultural chemicals can impair trophic linkages between OM-associated microbial communities and invertebrates.

## Ethics approval and consent to participate

Not applicable.

## Consent for publication

Not applicable.

## Availability of data and material

The datasets and/or scripts used for data analysis are available upon request.

## Competing interests

The authors state that they have no conflict of interest.

## Funding

The study did not receive external funding.

## Notes

### Competing Interest Statement

The authors have declared no competing interest.

